# Autophagy Restricts Tomato Fruit Ripening Via a General Role in Ethylene Repression

**DOI:** 10.1101/2023.12.20.572633

**Authors:** Girishkumar Kumaran, Pradeep Kumar Pathak, Ebenezer Quandoh, Sergey Mursalimov, Jyoti Devi, Sharon Alkalai-Tuvia, Jia Xuan Leong, Kyrylo Schenstnyi, Elena Levin, Suayib Üstün, Simon Michaeli

## Abstract

Autophagy, a cellular degradation pathway, and the phytohormone ethylene function in plant development, senescence, and stress responses. However, the manner of their interaction is mostly unknown. We reasoned that this may be revealed by studying autophagy in a climacteric fruit ripening context, for which ethylene is crucial. During ripening, fruits undergo softening, color change, toxic compound degradation, volatile production, and sugar assembly by fine-tuning synthesis and degradation of their cellular content. For autophagy activity assessment, we analyzed autophagy-related 8 (ATG8) lipidation and GFP-ATG8-labeled autophagosome flux in tomato fruit cells. Autophagy activity increased sharply from ripening initiation, climaxed at its middle stage, and declined towards its end, resembling ethylene production dynamics. Silencing the core-autophagy genes *SlATG2*, *SlATG7*, and *SlATG4* separately in mature fruits resulted in early ethylene production and ripening onset, which was abrogated by 1-methylcyclopropene (1-MCP), an ethylene signaling inhibitor. Beyond ripening, Arabidopsis *atg5* and *atg7* mutant seedlings exhibited elevated ethylene production and sensitivity to 1-Aminocyclopropane 1-carboxylic acid (ACC), ethylenès precursor, which induces autophagy. This research demonstrates that autophagy limits tomato fruit ripening via a general role in ethylene restriction, opening the path for a mechanistic understanding of autophagy-ethylene crosstalk and harnessing autophagy for fruit shelf-life extension.

## INTRODUCTION

Ripening is governed by myriad molecular mechanisms, including phytohormones, subsequent signaling pathways, epigenetic elements, and ubiquitin-proteasome system (UPS) proteolysis (Giovannoni et al., 2017). It involves cell wall, toxic compounds, and starch breakdown, contributing to fruit softening, edibility, and taste. An additional and pronounced phenomenon is the transition from green, none-ripe to colorful, ripe fruit that attracts humans and other animals. To allow these changes, fruits must constantly reshape their proteome, presumably by fine-tuning protein degradation and synthesis events (Szymanski et al., 2017).

In climacteric fruits, such as banana, mango, avocado, and tomato, ripening is accompanied by respiration and ethylene bursts (Cherian et al., 2014; Giovannoni et al., 2017; Huang et al., 2022; Osorio et al., 2013). Once harvested at a mature-green (MG) stage, this type of fruit will continue ripening postharvest. In tomato fruits, ethylene production increases upon entry into the breaker stage and peaks at around the orange/pink stages, after which a decline is detected towards the red-ripe stage (Fenn and Giovannoni, 2021; Osorio et al., 2013; Huang et al., 2022). While the UPS was shown to be critical in ethylene signaling and ripening, knowledge of the impact of autophagy, another central degradation system, is rather limited.

Autophagy delivers cytosolic components, such as organelles and macromolecules, to the vacuole for degradation and recycling. Double-membrane vesicles, termed autophagosomes, are generated around the cellular cargo destined for degradation. The autophagosome then fuses with the tonoplast to release a single membrane structure, termed autophagic-body. Inside the vacuole, the autophagic body degrades along with its cargo, and its constituents are recycled to replenish cellular energy. Autophagy is regulated and executed via the coordinated function of more than thirty autophagy-related (ATG) proteins (Marshall and Vierstra, 2018; Ding et al., 2018). Among these, the ubiquitin-like ATG8 proteins are important for autophagosome biogenesis, fusion with the vacuole and selective recognition of the cargo to be degraded. They are found either conjugated to phosphatidylethanolamine (PE) lipids (lipidated) or in a non-active free form (non-lipidated; Kellner et al., 2017). Members of the ATG8 conjugation machinery, such as ATG5 and ATG7, are classical targets for functional analysis in Arabidopsis as they are encoded by a single copy gene each. In tomato, there is a single *SlATG7* and two *SlATG5* genes, of which *SlATG5a* (Solyc02g036380) may be active (Zhou et al., 2014). ATG4, the protease that allows ATG8 lipidation or de-lipidation, is encoded by a single gene in tomato, while Arabidopsis harbors two members (Seo et al., 2016).

Initially, it was assumed that ethylene and autophagy are not linked based on the lack of leaf senescence recovery phenotype in Arabidopsis plants deficient in both autophagy and Ethylene-Insensitive 2 (*EIN2)* genes (*atg5ein2-1* or *atg2ein2-1* double mutants; Yoshimoto et al., 2009). However, several studies later suggested that crosstalk between ethylene and autophagy may exist (Liao et al., 2022). First, the transcript abundance of many ethylene biosynthesis and some ethylene-signaling genes was higher in *atg5* and *atg9* mutants than in WT (*Col-0*) Arabidopsis plants (Masclaux-Daubresse et al., 2014). Moreover, ACC-treated tomato plants exhibited increased autophagy activity and expression of SlATG8d and SlATG18h genes during drought. The authors suggested the direct binding of Ethylene Response Factor 5 (ERF5) to the promoters of these genes (Zhu et al., 2018). ACC was also reported to induce autophagy in Arabidopsis (Rodriguez et al., 2020). Finally, pollination-induced petal leaf senescence, accompanied by enhanced ethylene emission, was further accompanied by increased expression of petunia PhATG8 isoforms and generation of autophagosomes. Notably, 1-MCP delayed the induction of PhATG8 proteins following pollination and the appearance of the subcellular structures presumed to be autophagosomes (Shibuya et al., 2013). Nonetheless, direct evidence linking the two pathways is lacking.

To decipher the role of autophagy in tomato fruit ripening, we followed autophagy activity by monitoring SlATG8 lipidation and GFP-SlATG8 flux to fruit cell vacuoles. For functional analysis, we used two approaches that allow understanding of ripening-specific roles, avoiding possible pleiotropic effects of gene knockouts or constitutive knockdowns. First, we applied Virus-Induced Gene Silencing (VIGS) directly in MG fruits to silence *SlATG2* or *SlATG7*. Then, we generated *E8::SlATG4-RNAi* stable lines to silence *SlATG4* upon ripening induction. Under both methods, ripening was induced earlier than controls and was accompanied by increased ethylene production. Evaluation of Arabidopsis *atg5* and *atg7* seedlings showed their sensitivity to ethylenès precursor, ACC, and increased ethylene emission. Altogether, our results highlight a potentially ubiquitous function of autophagy in ethylene restriction, which confers a prime role in fruit ripening modulation.

## RESULTS

### Autophagy activity climaxes at mid-ripening

ATG8 family members are the proteins of choice for monitoring autophagy activity (Qi et al., 2023). Seven tomato proteins show high similarity to Arabidopsis AtATG8s, clustered into four subgroups (Fig. 1A), as was previously reported (Zess et al., 2019; Kellner et al., 2017). Data from the Sol Genomics Network - Tomato Expression Atlas (SGN-TEA; tea.solgenomics.net; Shinozaki et al., 2018) revealed that while the transcript level of five *SlATG8* genes remains relatively stable, the other two, *SlATG8-1.1* (Solyc08g078820) and *SlATG8-2.2* (Solyc07g064680), showed increasing expression along ripening progression (Fig. 1B). To verify that *SlATG8-2.2*, as a representative, encodes an autophagy-associated protein in fruit cells, we generated transgenic tomato plants expressing GFP-SlATG8-2.2. Confocal laser-scanning microscopy (CLSM) of fruit endocarp cells at the breaker stage highlighted small spherical bodies of about 1-2 µm in diameter reminiscent of autophagosomes (Fig. 1C, images C1-C3). These structures were motile (Movie S1), and occasionally seemed as ring-like structures. Shown here is a presumably mature phagophore at an advanced stage before closure (Fig. 1C, image C4, edges defined by arrowheads), later appearing as a fully closed and complete autophagosome (Fig. 1C, image C5).

**Fig. 1.**
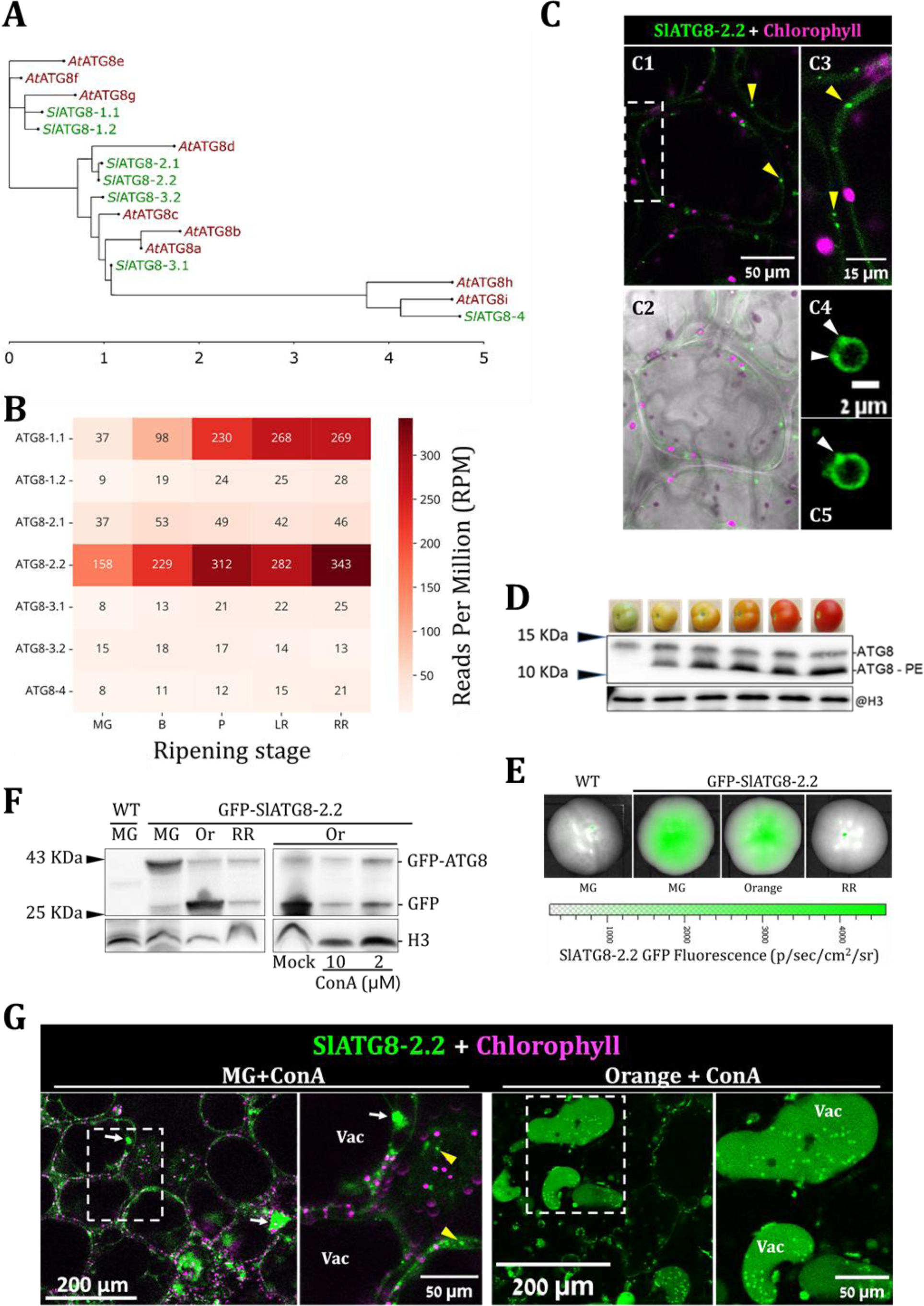
Autophagy activity increases along ripening progression. **A.** A phylogenetic tree showing the four tomato ATG8 (SlATG8) family subgroups with the related Arabidopsis AtATG8 members. The scale bar represents branch length in substitutions per site. **B.** Modified representation of *SlATG8* family members’ relative transcript expression in reads per million (RPM). Data obtained from SGN-TEA. **C.** CLSM imaging of fruit endocarp cells expressing GFP-SlATG8-2.2 (green). The Magenta signal represents chlorophyll autofluorescence. Yellow arrowheads point towards autophagosomes. Image C2 also includes the bright-field channel. Image C3 is an enlargement of the area defined with a dashed rectangle at C1. Images C4 and C5 show a developing autophagosome imaged one minute apart. White arrowheads in image C4 denote the estimated location of phagophore edges. The white arrowhead in image C5 points towards the estimated autophagosome closure site. **D.** Total pericarp proteins, from six ripening stages, were separated in resolving gel supplemented with 6M Urea and immunoblotted with @ATG8 antibodies (Abcam). Histone 3 (@H3; Millipore) serves as a loading control. A representative blot of three biological repeats is shown. **E.** Fluorescence emission from GFP-SlATG8-2.2 fruits at three ripening stages: mature-green (MG), orange, and red-ripe (RR), imaged using IVIS. None of the 22 RR-stage fruits that were examined showed any fluorescence. **F.** Total pericarp proteins of GFP-SlATG8-2.2 fruits from three ripening stages and WT MG-stage as control were immunoblotted with @GFP. Histone 3 (H3) serves as a loading control. Orange-stage (Or) fruits were also treated with the indicated concentrations of ConA or mock (DMSO). Representative blots of three biological repeats are shown. **G.** CLSM imaging of ConA-treated endocarp cells of MG- and orange-stage GFP-SlATG8-2.2 fruits. The Magenta signal represents chlorophyll autofluorescence. Yellow arrowheads point towards autophagosomes. White arrows denote the location of cytosolic aggregates, potentially culminating due to their inability to degrade. Representative images of 10 replicates (representing more than 50 cells) per ripening stage are shown. No autophagic bodies were detected in any of the MG-stage cells. Vac, vacuole lumen.

To gain insight into autophagy activity, we immunoblotted total pericarp protein extracts of tomato fruits at different ripening stages with an anti-ATG8 antibody. We suspect this polyclonal antibody recognizes all expressed ATG8 isoforms. Our analysis showed a sharp increase in ATG8-PE levels, from undetectable levels at the MG stage to prominent expression at mid-ripening, and maintaining these high levels till the final red-ripe stage (Fig. 1D). The increase in ATG8-PE may reflect either an elevation in autophagy activity or a blockage in autophagy flux to the vacuole. To determine which of these options is factual, GFP-SlATG8-2.2 expressing fruits were evaluated to follow autophagy flux. Using an In-Vivo Imaging System (IVIS; Perkin Elmer), we detected sufficient GFP fluorescence in fruits of early-to-mid ripening stages (MG to orange). However, the signal wasn’t detected in any of the red-ripe stage fruits that were examined (n=22; Fig. 1E). We then examined autophagy flux via the GFP-release assay by immunoblotting GFP on fruit protein extracts of three ripening stages, MG, orange, and red-ripe. Results showed a sharp increase in free-GFP level (i.e., increasing flux) from the MG to the orange stage (Fig. 1F). Consistent with our IVIS analysis (Fig. 1E), the level of GFP detection at the red-ripe stage was significantly lower than the earlier stages. Nonetheless, it was sufficient to observe a shift back towards balanced levels of GFP-ATG8-2.2 relative to free-GFP, indicating a reduction in autophagy flux (Fig. 1F, RR sample). We then examined the impact of concanamycin A (ConA), a drug that inhibits vacuolar degradation, on the flux at the orange stage. We tried two concentrations, mild (2 µM) and high (10 µM), since fruit vacuoles tend to be more acidic than non-fruit vacuoles (Gong et al., 2021). ConA reduced the free-GFP level at orange-stage cells under both concentrations (Fig. 1G, right blot), suggesting that most of it is produced via vacuolar degradation. For further confirmation, CLSM analysis of orange-stage vacuoles following ConA treatment showed punctate structures corresponding to autophagic bodies (Fig. 1G), exhibiting the typical random movement within the vacuole lumen (Movie S2). Notably, no such autophagic bodies were detected in any of the MG stage vacuoles following analysis of more than 50 cells in 10 different fruits (Fig. 1G). Altogether, increasing ATG8-PE levels along ripening renders increased autophagy activity, which peaks at the mid (orange) stage.

### Silencing key *ATG* genes in mature tomato fruits accelerates their ripening

Autophagy is involved in many processes, including energy homeostasis and nutrient remobilization (Avin-Wittenberg et al., 2015; Masclaux-Daubresse et al., 2017; Li et al., 2015). Therefore, knockout or constitutive knockdown of any key *ATG* gene would result in pleiotropic effects, preventing the understanding of ripening-specific roles. To overcome this, we applied two approaches. First, we used VIGS directly in MG-stage *Del/Ros1* tomato fruits, harboring the Snapdragon transgenes *Delila* (*DEL*) and *Rosea 1* (*ROS1*) driven by the E8 ripening-induced promoter. Therefore, silencing *Del/Ros1*, which prevents the ripening-associated appearance of purple fruits (Orzaez et al., 2009), is a visual indicator for silencing (Fig. 2A; see D/R vs. NT fruits). Silencing of the core autophagy genes, *SlATG2* or *SlATG7*, each with *DEL/ROS1* (*D/R/ATG2*-VIGS or *D/R/ATG7*-VIGS, respectively), resulted in SlATG8 accumulation, presumably due to its decreased turn-over (Fig. 2D). This is consistent with the accumulation of AtATG8 in Arabidopsis *atg2* and *atg7* mutants (Kang et al., 2018; Munch et al., 2014; Luo et al., 2023). Phenotypically, silencing *ATG2* resulted in accelerated color transition and softening (Fig. 2A and B). Quantitative examination of fruit color change, manifested by a declining hue parameter, demonstrated a similar outcome for *ATG7*-silenced fruits (Fig. 2C).

**Fig. 2.**
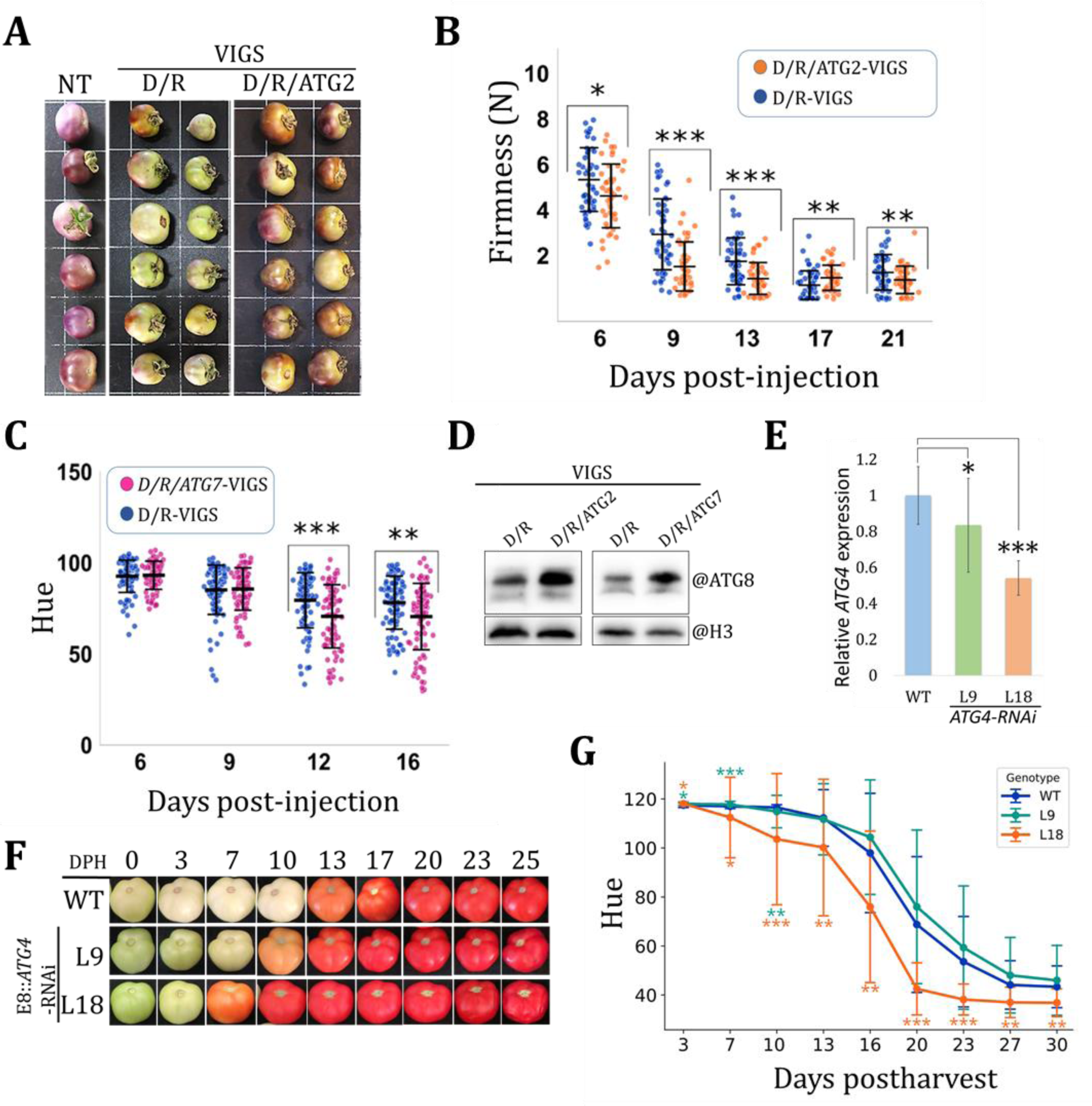
Silencing *SlATG2*, *SlATG7* or *SlATG4* in mature fruits results in accelerated ripening. **A-D.** D/R-VIGS, silencing of *Del*/*Ros1.* D/R/ATG2-VIGS, silencing of *Del*/*Ros1* and *ATG2*. D/R/ATG7-VIGS, silencing of *Del*/*Ros1* and *ATG7*. **A.** Representative *Del*/*Ros1* tomato fruits at 10 days post injection of agrobacterium cells harboring the indicated VIGS constructs. NT, non-treated. **B.** Firmness measurements of populations of VIGS-treated Fruits. Firmness is displayed as the force in Newton (N) required to achieve a similar probe penetration (see Materials and Methods) on different days post agrobacterium injection. Values are shown as the mean ± standard deviation (n > 30) **C.** Quantification of the green-to-red color transition (hue) of VIGS-treated fruits at the indicated days post agrobacterium injection. Decreasing hue values indicate color transition. Values are shown as the mean ± standard deviation (n = 30 with three technical measurements for each fruit). **D.** Total pericarp proteins from VIGS-treated fruits were immunoblotted with @ATG8 antibodies (Abcam). Histone 3 (@H3; Millipore) serves as a loading control. Representative blots of two biological repeats are shown. **E.** Relative expression of *SlATG4* in pink-stage fruits from WT and two E8::ATG4-RNAi lines was measured by quantitative RT-PCR. Values are shown as the mean ± standard error after normalization to three reference genes (TIP4, Actin, and GAPDH). Significance was tested relative to WT using one-tailed and paired *student t-tests;* n = 9. *, *0.01 < p < 0.05.* ***, *p < 0.001*. **F.** WT, L9, and L18 tomato fruits across 25 days postharvest. The first fruit to initiate ripening out of the sampled population of each genotype is shown here. **G.** Quantification of the green-to-red color transition (hue) of WT, L9, and L18 fruits at the indicated days postharvest. Values are shown as the mean ± standard deviation (n = 30). **B, C, and G.** Significance were tested relative to controls (D/R-VIGS in panels B and C, or WT in panel G) for each time point. Either unpaired and two-tailed *Student t-tests* or the *Mann-Whitney U* test were performed based on population distribution and variance (see Materials and Methods). No asterisk, not significant. *, *0.01 < p < 0.05*. **, *0.001 < p < 0.01*. ***, *p < 0.001*.

To validate the role of autophagy in a stable silencing system, we utilized an *SlATG4-RNAi* construct to silence the sole *ATG4* representative in the tomato genome. Similarly to *SlATG8-1.1* and *SlATG8-2.2*, *SlATG4* expression increases along ripening progression (Fig. S1), suggesting that ATG8 lipidation or delipidation may be important for ripening (Holla et al., 2023; Yoshimoto et al., 2004). Previously, it was reported that silencing *SlATG4* throughout plant development (using 35S::*SlATG4-RNAi* lines) results in early leaf senescence and reduced fruit yield (Alseekh et al., 2022). To examine ripening-specific functions, we generated transgenic lines harboring the same construct, albeit under the regulation of the E8 ripening-induced promoter (E8::*SlATG4-RNAi*). We focused on line #18 (L18), which exhibited significant silencing, and L9, which was revealed as a relatively weak line (Fig 2E). Fruits from WT and the transgenic lines were harvested at the MG stage and monitored postharvest for color transition (Fig. 2F). On average, L18 fruits started ripening seven days before WT and L9 fruits (Fig. 2G). Early ripening initiation was also observed in non-detached fruits of E8::*SlATG4-RNAi* lines (Fig. S2; L18 and L15 are presented). Altogether, these results demonstrate that autophagy is involved in ripening restriction.

### 1-MCP abrogates autophagy deficiency-induced ripening

To examine whether the impact of autophagy on ripening is linked to ethylene, we treated WT, L18, and L9 fruits with the ethylene signaling inhibitor, 1-MCP, before hue measurements were taken to assess color transition. Fruits were harvested at a relatively advanced stage of color transition (hue value of ∼105) to minimize the ripening initiation gap between WT, L9, and L18. Nonetheless, the latter still advanced faster than the others. Importantly, 1-MCP abolished the ripening advantage of L18 and exhibited almost identical delayed ripening dynamics as the WT fruits (Fig. 3A and B). This suggests that autophagy regulates ripening via an interaction with ethylene.

**Fig. 3.**
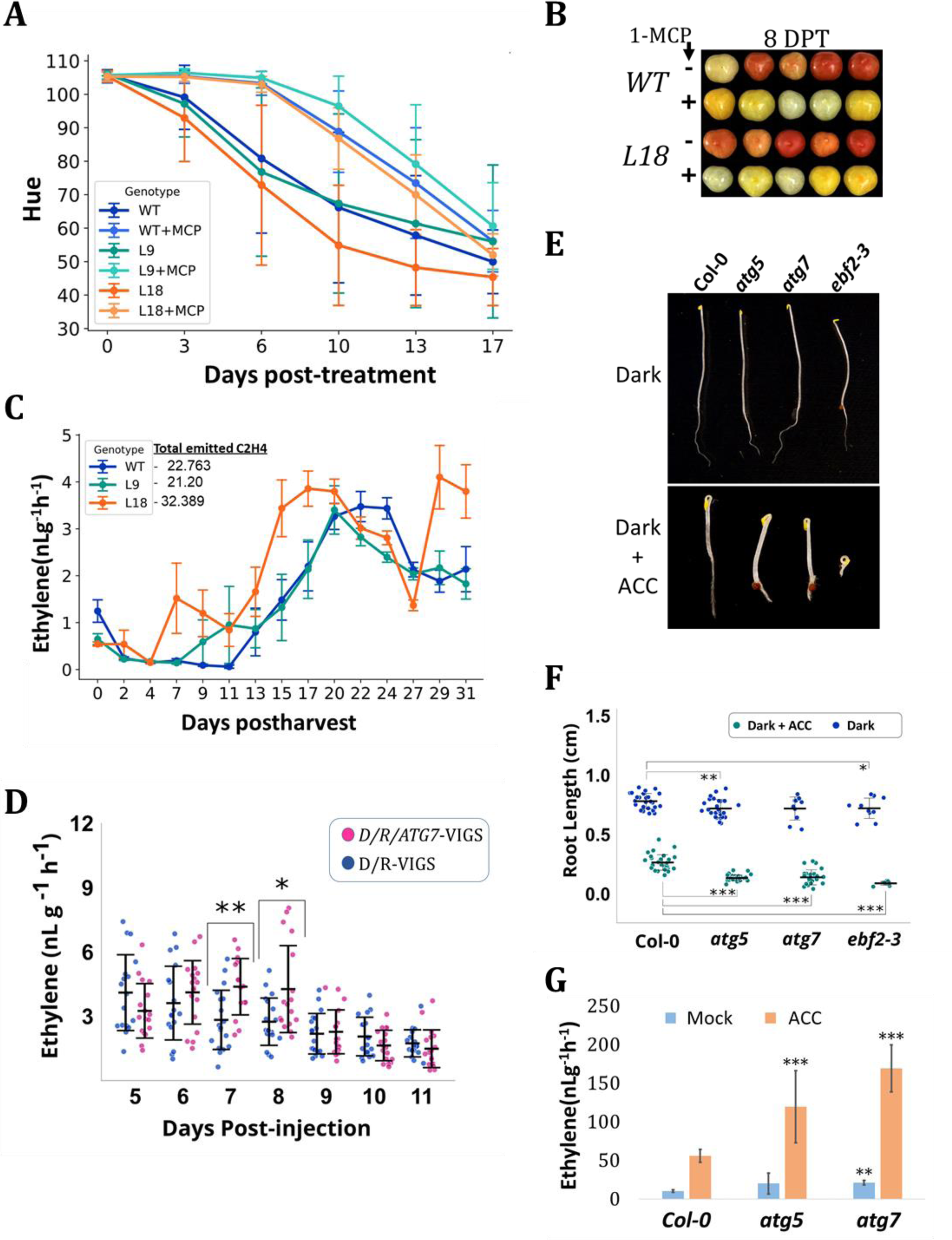
Autophagy deficiency in tomato fruits and Arabidopsis seedlings results in increased ethylene emission. **A.** Quantification of green-to-red color transition (hue) of WT, L9, and L18 fruits, treated or not with 1-MCP. Values at the indicated days post-treatment are shown as the mean ± standard deviation (n = 10). **B.** Representative WT and L18 fruits from the experiment described in panel A. **C.** Ethylene production in WT, L9, and L18 tomato fruits over 31 days postharvest. The numbers next to the legend represent total ethylene production, on average, for each genotype during the entire measurement duration. Values are mean ± standard error (n = 30). **D.** Ethylene production from Del/ROS1 fruits following *Del/Ros1* VIGS-mediated silencing (D/R-VIGS as control) or fruits additionally silenced in *ATG7* (D/R/ATG7-VIGS). Values at the indicated days post-injection of agrobacterium cells are shown as the mean ± standard deviation (n > 15; each measurement recorded from three pooled fruits). **E.** Representative Arabidopsis seedlings of the indicated genotypes assessed for ACC-induced triple-response assay (Dark-grown, etiolated seedlings, with or without ACC treatment; Material and Methods). **F.** Root length measurements of seedlings subjected to the triple-response assay. Values represent means ± standard deviation (n > 10). **G.** Ethylene production from 10 days old Arabidopsis seedlings. Values represent mean ± standard deviation (n = 5, each measurement recorded from 30 pooled seedlings). **D, F, and G.** Significance was tested relative to controls (D/R-VIGS in panel D or *Col-0* in panels F and G). Either unpaired and two-tailed *Student t-tests* or the *Mann-Whitney U* test were performed based on population distribution and variance (see Materials and Methods). No asterisk, not significant. *, *0.01 < p < 0.05*. **, *0.001 < p < 0.01*. ***, *p < 0.001*.

### Autophagy deficiency in tomato fruits and Arabidopsis seedlings results in increased ethylene emission

Examination of ethylene emission showed earlier climacteric phase onset and total higher ethylene production in L18 compared to WT and L9 fruits (Fig. 3C). To clarify whether this is a general autophagy function, we examined ethylene emission from the *Del/Ros1/ATG7-*VIGS fruits compared to the *Del/Ros1-*VIGS reference. Experiments showed significantly elevated ethylene emission at 7 and 8 days post injection (Fig. 3D). To see whether this impact of autophagy goes beyond ripening, we examined Arabidopsis *atg5-1* and *atg7-2* knockout seedlings under the triple-response assay, a known test for detecting ethylene-sensitive or insensitive mutants (Merchante and Stepanova, 2017). Results showed that, on average, *atg* mutants had shorter roots than WT seedlings under dark (significant only for *atg5*), which became more prominent following ACC application (Fig. 3E and F). *ebf2-3* was used as a known ethylene hypersensitive reference mutant (Guo and Ecker, 2003; Potuschak et al., 2003; Gagne et al., 2004). Notably, *atg5* and *atg7* seedlings exhibited a 2- and 3-fold increase in ethylene emission following ACC treatment, respectively. With *atg7* exhibiting elevated ethylene emission also under mock treatment (Fig. 3G). These results suggest that the triple-response phenotype of *atg* mutants is due to increased ethylene production, highlighting a potentially general role for autophagy in ethylene repression.

### ACC induces autophagy activity

It was previously reported that ACC induces autophagy based on the analysis of GFP-ATG8a foci numbers (without ConA), GFP-release assay, and expression of the selective-autophagy cargo-receptor, NBR1 (Rodriguez et al., 2020). Based on our observations, we wanted to reaffirm this with complementary approaches. First, we tested ATG8 levels in Arabidopsis Col-0 seedlings subjected to ACC or mock. Results showed that both total ATG8 (Fig. 4A; upper blot) and ATG8-PE (Fig. 4A; lower blot) levels were higher in ACC-treated seedlings at 72 and 96 h post-treatment. Then, we followed GFP-ATG8e in Arabidopsis roots subjected to combinations of ACC, ConA or mock (Fig. 4B). Quantification revealed increased amount of puncta whether ACC was applied solely or combined with ConA, demonstrating that ACC increases autophagosomes production and their flux to the vacuole (Fig. 4C). Together with the report by Rodriguez *et al*., these results confirm the ability of ACC to induce autophagy.

**Fig. 4.**
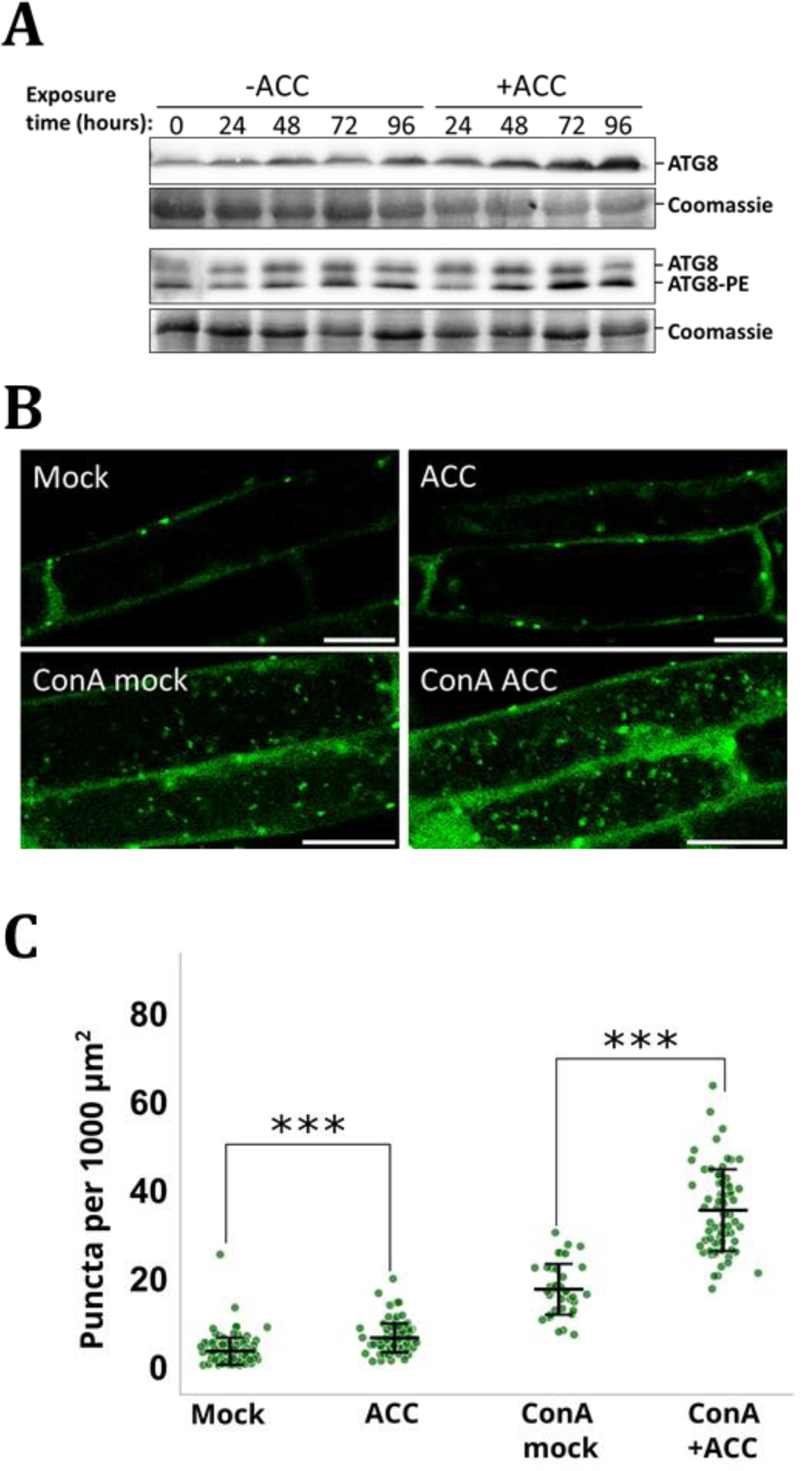
ACC induces autophagy. **A.** Total protein extracts of Arabidopsis seedlings, grown with or without ACC-containing media for the indicated time points (in hours), were immunoblotted with @ATG8. The upper lane shows standard separation, while the lower lane shows separation in a 6M urea-supplemented gel. Coomassie staining serves as loading control. **B.** Representative confocal images of Arabidopsis seedlings expressing GFP-ATG8e following treatment with ACC, mock, ConA + mock, or ConA + ACC. Scale-bars denote 20 µm **C.** Quantification of GFP fluorescence-emitting puncta in image slices of plants treated as described in panel B. Numbers were normalized to image area (1000 µm^2^). Significance was tested relative to respective controls (mock-treated samples). Either unpaired and two-tailed *Student t-tests* or the *Mann-Whitney U* test were performed based on population distribution and variance (see Materials and Methods). n > 35. ***, *p < 0.001*.

## DISCUSSION

Autophagy activity during ripening (Fig. 1) strikingly resembles the climacteric ethylene production trend (See WT in Fig. 3C, model in Fig. 5A and Wheeler et al., 2004; Cherian et al., 2014; Huang et al., 2022). This was the first hint for an ethylene-related role of autophagy during ripening. Indeed, results showed that silencing *SlATG4, SlATG2,* or *SlATG7* promotes ripening (Fig. 2) by allowing premature and increased ethylene production (Fig. 3). It is well-accepted that there is a physiological and molecular resemblance between ripening and senescence, as can be seen in dry fruits such as Arabidopsis siliques (Seymour et al., 2013; Gómez et al., 2014; Keren-Keiserman et al., 2022). Autophagy is an anti-aging mechanism in plants and animals (Aman et al., 2021; Liu and Bassham, 2012; Minina et al., 2018), reconciling with a repressive effect on ripening.

**Fig. 5.**
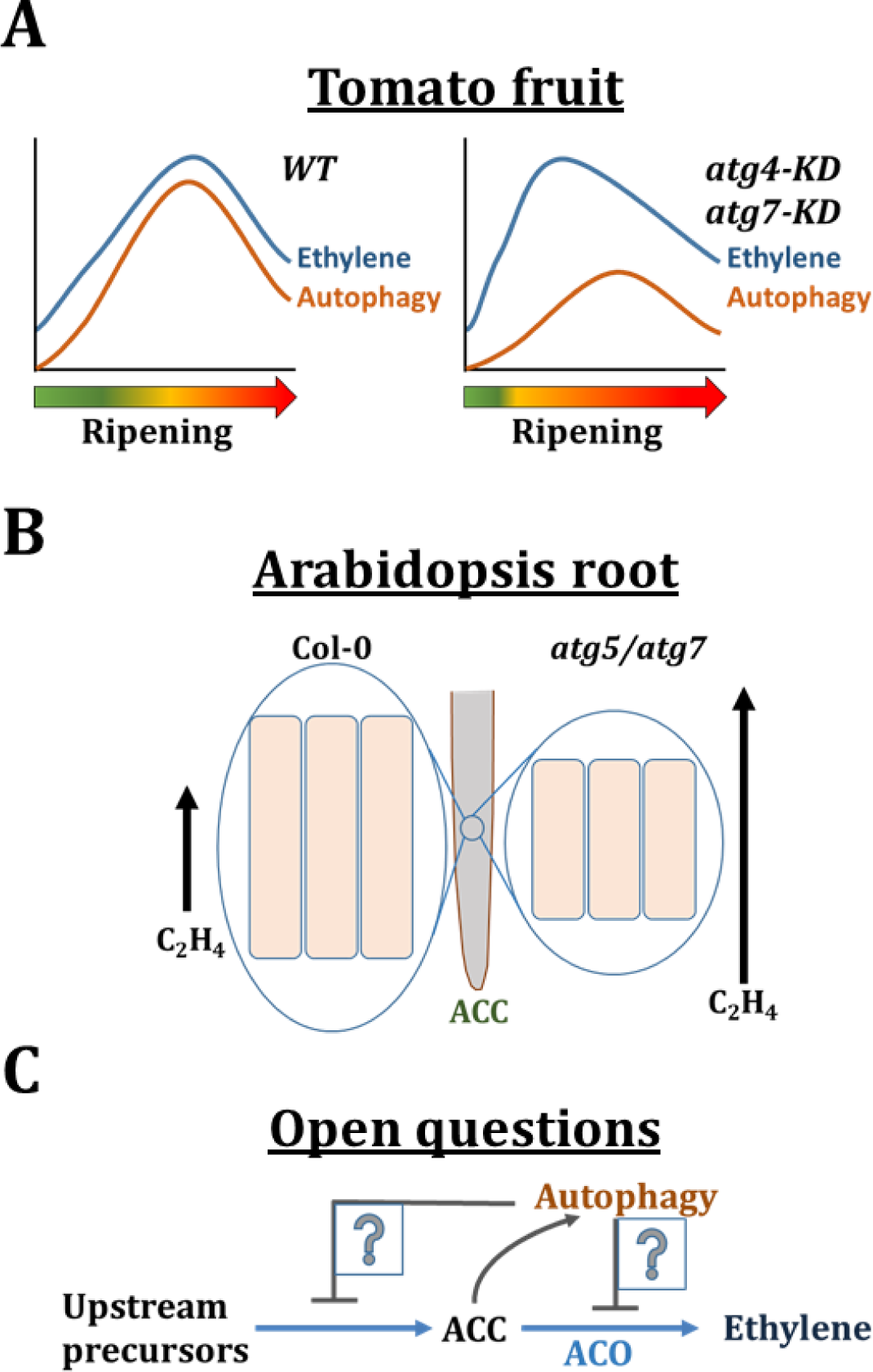
Current working model and open questions. **A.** Representation of ethylene and autophagy dynamics along WT or *ATG*-knockdown fruit ripening. In WT, autophagy and ethylene share similar dynamics, and autophagy participates in buffering ethylene levels. In *ATG* knockdown fruits, the milder buildup of autophagy results in earlier induction of climacteric ethylene production and, hence, earlier ripening. **B.** Representation of ethylene production in *Col-0* or *ATG-*deficient Arabidopsis seedlings and its impact on root elongation. Following ACC application, *atg5* or *atg7* mutants produce more ethylene, leading to reduced root cell elongation compared to Col-0. **C.** Autophagy may potentially restrict any of the steps leading to ethylene production. Moreover, ACC induces autophagy, raising the possibility of a feedback-loop interaction.

We propose that the milder buildup of autophagic capacity in the *SlATG* silenced fruits permits earlier ethylene production, leading to earlier ripening commencement (Fig. 5A). Intriguingly, the participation of autophagy in non-climacteric fruit ripening of pepper and strawberry was recently reported (López-Vidal et al., 2020; Sánchez-Sevilla et al., 2021). Contrary to our results, it was concluded that autophagy acts in strawberry ripening promotion (Sánchez-Sevilla et al., 2021). If so, we can’t rule out the possibility of autophagy having contradicting roles in climacteric and non-climacteric fruits. An intriguing possibility is that this reflects the different impact of ethylene on both fruit types (Cherian et al., 2014; Osorio et al., 2013).

The increased sensitivity of Arabidopsis *atg5* and *atg7* mutants to ACC, coupled with their elevated ethylene production (Fig. 3E-G), suggest that the repressive effect of autophagy is potentially widespread and may extend to other roles of ethylene beyond ripening. For example, during root cell elongation (Fig. 5B).

Several questions emerge from this study. First, how does autophagy limit ethylene? This may be happening via the selective degradation of components involved in ethylene or ACC production, such as ACC-Synthase (ACS) or ACC-Oxidase (ACO) enzymes (Fig. 5C; Houben and Van de Poel, 2019; Park et al., 2021). Elimination of even further upstream precursors or other regulatory elements of ethylene production is another possibility. Finally, it can’t be excluded that autophagy may also regulate ethylene signaling components (Binder, 2020). Notably, the ability of ACC to induce autophagy highlights a potential feedback loop for ethylene regulation (Fig. 5C). Additional questions are: What is the weight of ethylene in the well-known early senescence phenotype of autophagy mutants? Especially since, so far, it has mostly been associated with salicylic acid (Yoshimoto et al., 2009). Finally, does autophagy regulate ripening solely via its impact on ethylene? Our results don’t suggest that ethylene modulation is the only function of autophagy during fruit ripening, but they do suggest it is central. Although considerable reprogramming of fruit proteome can be attributed to the UPS, it is reasonable to assume that other components, such as protein complexes, other macromolecules, and organelles, are recycled via several degradation pathways, including selective-autophagy (Clavel and Dagdas, 2021). Further studies are required to settle these questions.

## MATERIALS AND METHODS

### Plants growth and transformation

Tomato Micro-Tom and Moneymaker cultivars and Arabidopsis Col-0 ecotype were used in this study. Tomatoes were grown in pots at 23°C in a climate-controlled greenhouse under a 12 h light/dark cycle. Arabidopsis was grown in growth chambers at 22°C and 16 h light/8 h dark cycles. Transgenic Del/Ros1 Micro-Tom (Orzaez et al., 2009) seeds were obtained from Prof. Asaph Aharoni’s lab (Weizmann Institute). GFP-SlATG8-2.2 Moneymaker lines were generated by transforming the 35S::GFP-StATG8-2.2 construct (Potato StATG8-2.2 is identical to the tomato counterpart), which was kindly provided by Dr. Yasin Dagdas (Zess et al., 2019). E8::SlATG4-RNAi lines were generated in Micro-Tom background.

Tomato transformation was done with GV3101 agrobacteria harboring the appropriate plasmids as described in (Sun et al., 2015) with minor modifications. Tomato seeds were surface sterilized and germinated on 1/2 MS (+ 30 g/L sucrose) media. After ten days, fully expanded cotyledons were dissected, with only their central sections used. Explants were immersed in Agrobacterium suspensions (OD600 = 0.6) for 20 minutes. Explants were then gently dried on Whatman paper and transferred to a co-cultivation medium (MS medium supplemented with 3% (w/v) sucrose, 0.8% Agar, and 2 mg/L BAP) for 2 days in the dark. Then explants were transitioned to a shoot induction medium (MS medium supplemented with 1 mg/L Zeatin, along with 0.8% (w/v) Agar, 3% (w/v) sucrose, 50 mg/L kanamycin, and 500 mg/L cefotaxime). Incubation took place at 26°C with a 16-hour photoperiod. After 30 days, plantlets were separated from the original explants and transferred to a fresh induction media. Shoots reaching lengths of 2–3 cm were excised and transplanted to a root induction medium (MS medium containing 1 mg/L IAA, 0.3% (w/v) Agar, 3% (w/v) sucrose, and 50 mg/L kanamycin). All substances used were from Duchefa Biochemie.

GFP-SlATG8-2.2 Moneymaker line was generated similarly with a few differences. The seeds were sown on the germination medium (4.3 g/L MS including vitamins, 30 g/L sucrose, 100 mg/L Myo-inositol, 0.8% phytoagar, pH = 5.8). 2 μL of the Agrobacterium suspension (10 mm MgSO4, 200 μm acetosyringone; OD600 = 1.0) were applied per cotyledon and they were co-cultivated on the germination medium supplemented with 1 mg/L BAP and 1 mg/L NAA for 2 days in the dark at 22°C. In two days, the cotyledons were placed abaxial surface down on the germination medium supplemented with 35 mg/L kanamycin, 1 mg/L trans-Zeatin, and 250 mg/L ticarcillin disodium/clavulanate potassium. The cotyledons were placed in 14 h light/10 h dark, 23°C, 50% humidity, and were transferred to a fresh selection medium every seven days. In the second and fourth weeks, kanamycin concentration was increased to 50 mg/L and 100 mg/L, respectively. Regenerating shoots were cut at the base and transferred to the germination medium supplemented with 20 mg/L kanamycin, 0.1 mg/L IAA, and Vancomycin (500 mg/L).

### Plasmids construction

We employed the ClonExpress II One Step Cloning (Vazyme Biotech) and the Gateway Cloning system (Thermofisher Scientific) for VIGS-related cloning. The VIGS tool at the Sol Genomics Network website (vigs.solgenomics.net) was used to select appropriate 300 base-pair sequences. The primers used are listed in Table 1. ∼300 bp fragments of *SlATG2* (Solyc01g108160) and *SlATG7* (Solyc11g068930) were PCR amplified and cloned into pENTER-Gus (Thermofisher Scientific; modified to have spectinomycin, instead of kanamycin, resistance). Then, the Gateway LR reaction was performed to allow the transfer of the gene fragment into the pTRV2-Del/Ros1 vector. For ripening-specific *SlATG4* (Solyc01g006230) silencing, the E8::SlATG4-RNAi vector was generated by switching the promoter in the 35S::SlATG4-RNAi expression cassette (Alseekh et al., 2022). The original plasmid was amplified (Phusion polymerase) as a linear fragment without the 35S part. Then, ClonExpress II was employed to introduce a PCR-amplified E8 promoter with compatible flanking regions of the linearized plasmid. All cloned plasmids were transformed into the Mix & Go DH5alpha competent cells (Zymo Research, USA) and selected on LB agar plates containing the relevant antibiotics. Colony PCR confirmed positive clones, and plasmids were then purified using the Promega plasmid purification kit.

**Table 1:**
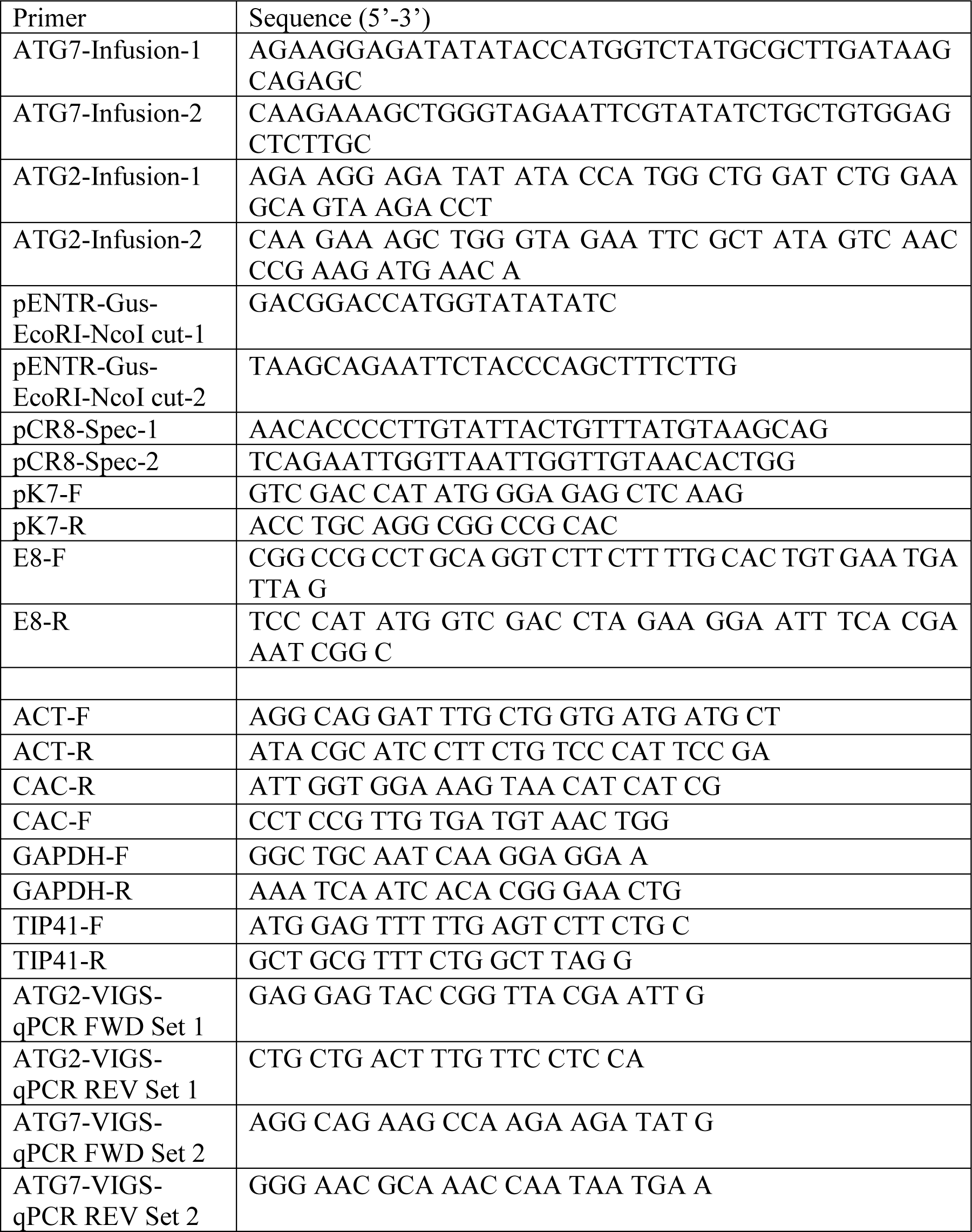
Primers Used in This Study.

### Immunoblotting

Liquid N frozen tomato fruit pericarp samples were manually ground to a fine powder using a mortar and pestle. 1 g samples were mixed with 500 µL of a 1x Laemmli buffer. The mixture was then heated to 95°C for 5 minutes and centrifuged at 16,000 x g (4°C) for 10 minutes. The resulting supernatant was collected for immunoblotting. To allow the separation of ATG8 and ATG8-PE fragments, equal amounts of protein were loaded onto custom-made 15% SDS-PAGE gels supplemented with 6M urea in the resolving gel. Gels were blotted onto PVDF membranes using a Bio-Rad transfer apparatus. The blots were incubated overnight at 4°C with 1:1,000 anti-ATG8 antibodies (Abcam) and subsequently with goat-anti-rabbit-HRP (1:10000; Jackson ImmunoResearch) before visualization in the Fusion Plus Imaging System (Vilber). Anti-H3 antibodies (EMD Millipore Corp.) were used for loading visualization (1:10,000). For the GFP-release assay, total protein extracts were loaded onto commercial precast gels (SurePage; GenScript) and immunoblotted with anti-GFP antibodies (Abcam Ab290; 1:2000). Arabidopsis samples were processed as previously described (Michaeli et al., 2019).

### RNA isolation and qRT-PCR

Total RNA was extracted from pink-stage tomato fruits using the Plant Spectrum RNA isolation kit. RNA concentration and purity were determined, and cDNA was synthesized from 1 µg of total RNA using the verso cDNA synthesis kit (Thermo Fisher). For qPCR, the Luna® Universal qPCR Master Mix (New England Biolabs) was used, and samples were processed in the StepOnePlus Real-Time PCR System (Thermo Fisher). The comparative Cт (ΔΔCт) method was employed for data analysis, normalizing target gene expression to reference genes (TIP41, GAPDH, and Actin) and comparing the relative expression levels between samples.

### Virus-Induced Gene Silencing (VIGS)

pTRV1 and pTRV2-Del/Ros-1 harboring the SlATG7- or SlATG2-derived fragments were introduced into Agrobacterium strain GV3101 using electroporation (BIO-RAD). A 5 mL culture was grown overnight at 28°C in a medium containing kanamycin. The next day, the culture was transferred to a 50 mL LB medium containing antibiotics, 10 mm MES, and 20 µM acetosyringone and grown overnight in a 28°C shaker. Agrobacterium cells were harvested, resuspended in infiltration media (10 mm MgCl2, 10 mm MES, 200 µM acetosyringone), left at room temperature for 3 hours, and adjusted to an optical density (OD600) of 0.1. Agrobacterium cells were then infiltrated, using a 1 mL syringe, through the fruit’s peduncle into MG-stage fruits while they were attached to the plant.

### Fluorescence Imaging and Quantification of transgenic fruits using the In-Vivo Imaging System (IVIS)

GFP-SlATG8-2.2 tomato fruits were harvested at different ripening stages (MG, turning, and Red Ripe) and kept dark for 12 hours before evaluation. GFP fluorescence (465 nm excitation and 520 nm detection) was acquired and analyzed using IVIS Lumina II equipped with an XFOV-24 lens and Living Image 4.3.1 software (PerkinElmer, Waltham, MA, USA). Optical fluorescence is presented as radiance intensity (photons s^−1^ cm^−2^ steradian^−1^).

### Confocal Microscopy, ConA treatment and quantification

Confocal imaging was done using the Leica SP8 CLSM system at the Volcani Institute microscopy core facility. Thin sections of tomato fruit pericarp (using razor blades) or intact Arabidopsis seedlings were positioned between a microscope slide and a coverslip containing 1/2 MS liquid media. Excitation/detection-range parameters for GFP were 488 nm/500–550 nm, respectively, and emissions were collected using the system’s hybrid (Hyd) detectors. X20 dry (NA 0.75) or X63 water immersion (NA 1.3) objectives were used. Scanning was routinely done in “line” mode. The images were processed and analyzed using FIJI (ImageJ).

For ConA treatment of fruit cells, thin fruit sections were incubated in liquid 1/2MS supplemented with 10 µM ConA, or the corresponding amount of DMSO (mock), and incubated 7 hours before imaging or 15 hours for immunoblots. For Arabidopsis analysis, GFP-ATG8E seeds were germinated in liquid 1/2 MS (Duchefa Biochemie, Netherlands) medium containing 1% sucrose. 4-day-old seedlings were transferred into the same medium containing either 0.4% DMSO (Sigma-Aldrich; as mock), 20 µM ACC, 1 µM ConA (Santa Cruz), and 0.4% DMSO, or 1 µM ConA and 20 µM ACC. Seedlings were incubated for two hours in the dark and transferred to slides in corresponding media. Counting of labeled autophagosomes was performed manually on single confocal-slice images. Forty-five seedlings and 677 cells were analyzed in three independent experiments.

### Triple response assay

The triple response assay was done as described (Merchante and Stepanova, 2017) with minor modifications. Sterilized seeds were placed onto plates containing 1/2 MS media (Duchefa Biochemie) and 1% sucrose, with or without 20 µM ACC. The plates were then moved to 4° C in darkness for 48 hours before exposure to light for 2 hours. Then, the plates were relocated to the growth room (22°C) and kept in darkness for 72 hours.

### Phylogenetic Tree Construction

The amino-acid sequences of tomato (*Solanum lycopersicum*) and *Arabidopsis thaliana* ATG8 proteins were retrieved from Sol Genomics Network (solgenomics.net) and TAIR (www.arabidopsis.org) databases. Sequences were aligned using ClustalW in the EMBL-EBI web server (https://www.ebi.ac.uk/Tools/msa/clustalw2/) using the default settings. The resulting file was uploaded to IQ-TREE v1.6 (http://www.iqtree.org/) for phylogenetic analysis, which included ModelFinder, tree reconstruction, and ultrafast bootstrap with 1000 replicates. ModelFinder was used to determine the best-fit model, with the Bayesian Information Criterion (BIC) guiding the model selection, ultimately choosing LG+G4. This model, incorporating a gamma distribution with four rate categories, was used for managing rate heterogeneity among sites. Branch support values were calculated using the SH-aLRT test and ultrafast bootstrap approximation with 1000 replicates. Finally, the tree was visualized in Newick format, using the Interactive Tree of Life (iTOL) web-based tool (https://itol.embl.de/). The tree image was exported and refined using Inkscape to enhance clarity and presentation.

### Fruits Color (hue) and firmness measurements

Color values a* and b* and the hue parameter were measured using the Konica Minolta Chroma Meter CR-400 Series Version 1.11. Fruits were considered at the MG stage when -0.59 < a*/b* < -0.47 (Batu, 2004). Then, fruits were harvested, cleaned, and measured for hue and firmness on different days postharvest, as indicated in related figures. In the VIGS experiments, hue was measured at three different regions of each fruit. Firmness was measured using TA.XT Plus Texture Analyzer (Stable Microsystems, Surrey, UK). The probe had a diameter of 3 mm, and it penetrated the fruit to 5% of its diameter at a speed of 1 mm s−1, recording the maximal endpoint force in Newton (N).

### Ethylene production measurements and 1-MCP treatment

Harvested fruits were placed in 200 ml sealed containers for two hours. Ethylene was measured by injecting 10 mL of head-space air into a gas chromatograph (GC; Varian 3300, Walnut Creek, CA, USA) with a Flame ionization detector (FID) and an alumina column. Ethylene production was calculated as nanoliters produced per fruit fresh weight per hour (nLg^-1^h^-1^). For Arabidopsis, seedlings of the different genotypes were grown on 1/2 MS + 1% sucrose solid media. ACC and mock treatments were applied on ten-day-old seedlings by adding 20 µM ACC (dissolved in liquid 1/2 MS media) or mock (ACC-free 1/2 MS media) to the plates. Following 96 hours of incubation, the seedlings were removed from the plates, their weights were recorded, and they were then quickly sealed in 20 ml syringes containing 1 ml 1/2 MS liquid media, with or without ACC, for two hours. Each syringe contained 30 seedlings (with 5-10 syringes per genotype and treatment in each experiment). Head-space was then collected with a fresh 10 ml syringe (by penetrating the needle into the 20 ml syringe) and injected into the GC apparatus. For 1-MCP treatment, fruits from WT and the E8::ATG4-RNAi lines were placed in a sealed flask containing 600 ppb of 1-MCP (RIMI, Israel) for 20 h. Mock-treated fruits were placed in similar flasks containing air with no additives. Then, fruits were released and placed in a “shelf-life” chamber (22° C) for the indicated duration for hue measurements.

### Statistics

Statistical analysis and data visualization were conducted using Python 3.9.6 with the following libraries:

- NumPy 1.25.2 for numerical operations and data handling
- Pandas 2.0.3 for data importation and data frame handling
- Matplotlib 3.7.2 for basic plotting
- Seaborn 0.12.2 for advanced data visualization

Calculations were performed using the subpackage scipy.*stats* from the open-source SciPy python-based library. The Shapiro-Wilk test (scipy.stats.shapiro — SciPy v1.11.2) was used for normality determination, where *p-value* > 0.05 suggests the data are normally distributed. The Levene test (scipy.stats.levene) was used for equal variance determination where *p-value* > 0.05 suggests normal distribution. If two datasets showed equal variance and normal distribution, then an independent two-sample t-test (scipy.stats.ttest_ind) was used. Otherwise, a non-parametric Mann-Whitney U test (scipy.stats.mannwhitneyu) was used.

## SUPPORTING MATERIAL

**Fig. S1:** *SlATG4* expression increases along tomato fruit ripening progression.

**Fig. S2:** E8::ATG4-RNAi fruits display earlier ripening when attached to plants.

**Movie. S1:** GFP-ATG8-2.2 labeled autophagosomes movement within fruit endocarp cells

**Movie. S2:** GFP-ATG8-2.2 labeled autophagic bodies movement within ConA-treated endocarp cell vacuoles of an orange-stage fruit.

## ACKNOWLEDGMENTS

We thank Dr. Tamar Avin-Wittenberg (Hebrew University) for providing the 35S::ATG4-RNAi plasmid, Dr. Yasin Dagdas (Gregor Mendel Institute, Vienna) for sharing the 35S::GFP-StATG8-2.2 plasmid, Prof. Asaph Aharoni (Weizmann Institute) for providing the Del/Ros1 tomato seeds and the related pRTV2 vector, Dr. Yana Kazachkova for her assistance in establishing the fruit VIGS system in our lab, and Prof. Elazar Fallik (Volcani Institute) for providing tomato fruits for some of our initial experiments. We also appreciate the scientific discussions held with all who are mentioned here and the helpful comment of Prof. Jim Giovannoni (Boyce Thompson Institute, USA) during a scientific meeting.

## AUTHOR CONTRIBUTIONS

GK, PKP, EQ, S. Mursalimov, JD, SAT, EL, and S. Michaeli designed experiments.

GK, PKP, EQ, S. Mursalimov, JD, SAT, and EL conducted experiments. GK performed all VIGS experiments, data and statistical analysis, phylogenetic tree construction, and autophagy activity-related assays. PKP generated the E8:ATG4-RNAi tomato lines and performed most Arabidopsis experiments. EQ assisted in almost every aspect of the study and conducted hue, firmness, ethylene measurements, and GFP-release assay blots. S. Mursalimov performed confocal microscopy and quantification in Arabidopsis. JD examined the E8:ATG4-RNAi lines. SAT assisted in hue, firmness, and ethylene measurements and performed the 1-MCP experiments. EL performed qPCR.

JXL and KS generated the GFP-ATG8-2.2 tomato lines.

SÜ supervised students and contributed to the draft.

All authors discussed the results and contributed to editing the manuscript.

S. Michaeli conceived the study, supervised the work, and wrote the manuscript.

## FUNDING

This research was supported by the US-Israel Binational Agricultural Research & Development fund (BARD) grant IS-5553-22, and the Israeli Ministry of Agriculture and Rural Development research grant 20-06-0018 to S. Michaeli; ARO Postdoctoral Fellowship to PKP; Fellowship from the International School of the Faculty of Agriculture, HUJI, to EQ; Emmy Noether Fellowship GZ: UE188/2-1 from the Deutsche Forschungsgemeinschaft (DFG) to SÜ.

---

**Fig. S1.**
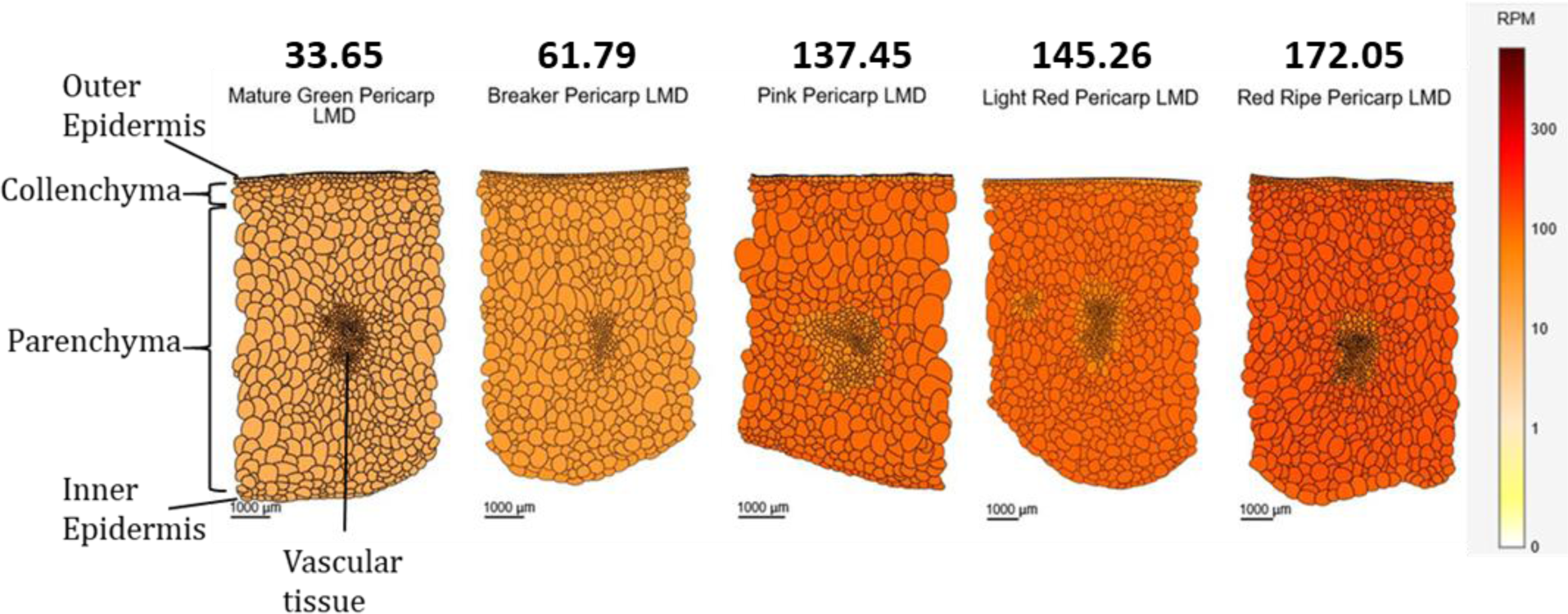
*SlATG4* expression increases along tomato fruit ripening progression. *SlATG4* (Solyc01g006230) expression in pericarp tissues, as retrieved following laser micro-dissection (LMD) and presented as reads per million (RPM) at the SOL genomics fruit expression atlas (SGN-TEA). Numbers above tissue illustrations indicate *SlATG4* RPM in the parenchyma cells.

**Fig. S2.**
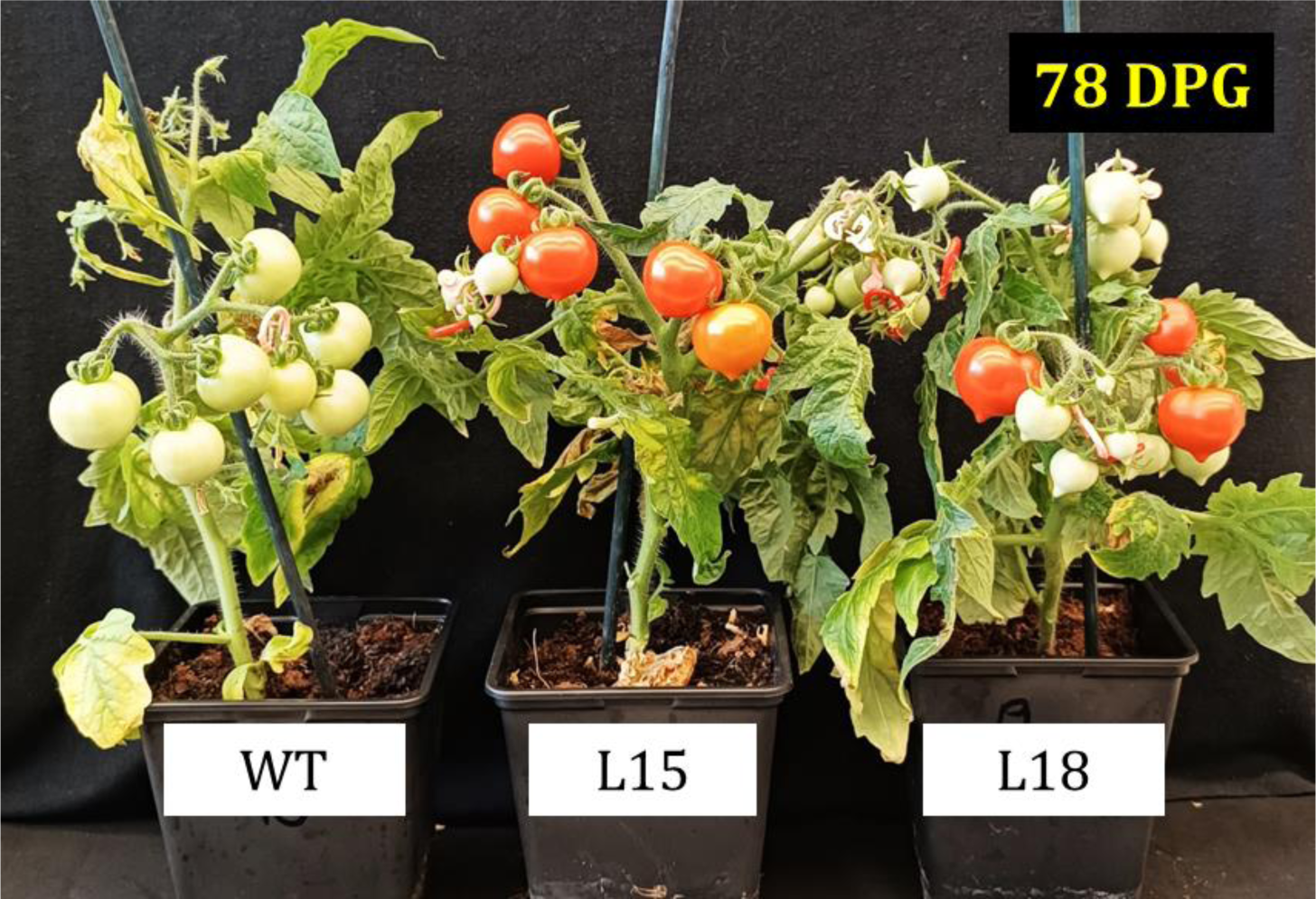
E8::ATG4-RNAi fruits display earlier ripening when attached to plants. Representative plants of WT and two E8::ATG4-RNAi lines at 78 days post-germination. Flowers of the three genotypes underwent simultaneous anthesis. Fruits of both L15 and L18 lines consistently ripened earlier than WT fruits in 4 independent growth cycles conducted in our greenhouse between March and October 2023.

## Notes

### Competing Interest Statement

The authors have declared no competing interest.

### Summary of Updates

The authors' affiliations were updated, and a few typos were corrected. The open questions model (Fig. 5C) was slightly revised.

